# Intraspecific variation in elemental accumulation and its association with salt tolerance in *Paspalum vaginatum*

**DOI:** 10.1101/2021.03.04.433795

**Authors:** David M. Goad, Elizabeth A. Kellogg, Ivan Baxter, Kenneth M. Olsen

**Affiliations:** Donald Danforth Plant Science Center, St. Louis, Missouri 63132; Department of Biology, Washington University in St. Louis, St. Louis, Missouri 63130

**Keywords:** *Paspalum vaginatum*, population structure, halophyte, ionomics, salt tolerance

## Abstract

Most plant species, including most crops, perform poorly in salt-affected soils because high sodium levels are cytotoxic and can disrupt uptake of water and important nutrients. Halophytes are species that have evolved adaptations to overcome these challenges and may be a useful source of knowledge for salt tolerance mechanisms and genes that may be transferable to crop species. The salt content of saline habitats can vary dramatically by location, providing ample opportunity for different populations of halophytic species to adapt to their local salt concentrations; however, the extent of this variation, and the physiology and polymorphisms that drive it, remain poorly understood. Differential accumulation of inorganic elements between genotypes or populations may play an important role in local salinity adaptation. To test this, we investigated the relationships between population structure, tissue ion concentrations (i.e., ionomic profiles) and salt tolerance in 17 “fine-textured” genotypes of the halophytic turfgrass seashore paspalum (*Paspalum vaginatum* Swartz). A high-throughput ionomics pipeline was used to quantify the shoot concentration of 18 inorganic elements across three salinity treatments. We found a significant relationship between population structure and ion accumulation, with strong correlations between principal components derived from genetic and ionomic data. Additionally, genotypes with higher salt tolerance accumulated more K and Fe and less Ca than less tolerant genotypes. Together these results indicate that differences in ion accumulation between *P. vaginatum* populations may reflect locally adapted salt stress responses.

## INTRODUCTION

High salt concentrations represent a particularly harsh environment for most terrestrial plants (Parida and Das 2005). As such, only a relatively few specialist species, called halophytes, have evolved to tolerate them. The challenge that wild plant species face in adapting to saline environments in the wild is also apparent in crop breeding, where the production of salt tolerant varieties in otherwise non-tolerant (i.e. glycophytic) crop species has proceeded slowly. This is problematic because 25%-30% of irrigated land is estimated to be salt-affected (Shahid et al. 2018) with an additional 10 million hectares becoming salinized each year (Szabolcs 1989). To develop crops that can produce satisfactory yields in these soils, a better understanding of how halophytes have managed to overcome extreme salt stress may be vital (Cheeseman 2015).

Currently, most of our knowledge of the genetic and physiological underpinnings of salt tolerance comes from crop species and plant model species that are not salt tolerant, with limited studies of halophytes. Furthermore, even studies in halophytes have often ignored intraspecific variation in levels of salt tolerance (Cheeseman 2013; Saslis-Lagoudakis et al. 2014). Some halophytic species grow in a wide range of salt concentrations, from brackish estuaries to beaches where they are regularly exposed undiluted seawater. If variation in salt tolerance is heritable, it could allow populations to adapt to the salt concentrations specific to their local environment. This sort of fine-scale adaptation could be immensely useful for not only incrementally increasing the tolerance of crop varieties, but also closely matching them to regional salinity conditions.

Ionomics, the quantification of the total inorganic contents of a plant, is a cost-effective and high-throughput method for studying how plants respond to stress (Baxter et al. 2008). Specifically, it can assess differences in the concentration of each element individually or by incorporating the concentration of multiple elements into an “elemental profile” for each plant. It is particularly useful for investigating intraspecific variation in salt tolerance because plants must deal with high external salt concentrations by controlling the uptake of Na while maintaining the accumulation of other important elements (Kumari et al. 2015). Sodium accumulation can be beneficial for osmoregulation in high salt environments; however, at high concentrations it is toxic. Additionally, Na can compete for uptake with other important elements such as Mg, K, and Ca, causing nutrient deficiencies (Yermiyahu et al. 1994; Ali and Yun 2017). By measuring differences in the concentrations of these and other elements in tissues between different genotypes, ionomics can provide a window into intraspecific variation in tolerance. For example, increased Na accumulation due to a polymorphism in a sodium transporter gene is associated with adaptation to saline environments in the glycophyte *Arabidopsis thaliana* (Baxter et al. 2010), and differences in the accumulation of several different ions have been reported between halophytic and glycophytic *Lotus* species (Sanchez et al. 2011). Studies examining ionomic variation within a halophytic species, however, remain rare.

The model halophyte seashore paspalum (*Paspalum vaginatum* Swartz) is an ideal model system to ask questions about intraspecific variation in salt tolerance. As a turfgrass with a dedicated breeding program and reference genome, it has been the focus of several studies which documented variation in salt tolerance between breeding lines (Lee et al. 2004a, 2004b, 2005, 2007, 2008). However, these studies have focused on breeding lines that had been collected for their favorable turf qualities and may inadvertently have excluded less salt-tolerant genotypes, thereby reducing the potential to detect phenotypic variation. Additionally, these studies predate genetic characterizations of the study lines, which subsequently revealed extensive genotypic redundancy and potential mislabeling in publicly available germplasm collections (Eudy et al. 2017; Goad et al. 2021). The lack of genotypic data in earlier studies also prevented comparisons between genetic populations.

While differences in salt tolerance between genotypes of *P. vaginatum* have been recorded, the mechanisms underlying this variation are not well understood. Unlike some halophytes, *P. vaginatum* does not seem to secrete salt and therefore must deal with sodium toxicity by either excluding sodium ions at uptake or sequestering them in specialized structures (Chen et al. 2009). Recent work has shown that epidermal leaf papillae may act as Na sinks in *P. vaginatum*, supporting the hypothesis of Na sequestration (Spiekerman and Devos 2020). Notably, these papillae are poorly developed in the closely related glycophyte *P. distichum.* Another potential salt tolerance mechanism is the use of organic osmolytes or inorganic ions for osmoregulation. There is some evidence of this mechanism as well. *Paspalum vaginatum* accessions with increased proline accumulation showed less reduction in biomass at increased salt concentrations (Lee et al. 2008), and K accumulation was associated with higher overall biomass at all salinity levels, but not with salt tolerance as indicated by the relative growth rate between low and high salt levels (Lee et al. 2007).

*Paspalum vaginatum* falls into two morphotypes, a “fine-textured” form which is sought out for turf and a “coarse-textured” form which is not. In a recent study Goad et al. (2021) showed that the fine-textured plants, including those from breeding populations at the US Department of Agriculture (USDA), are uniformly diploid and are genetically distinct from coarse-type accessions, which consist of interspecific hybrids of varying ploidy. That study also uncovered circumstantial evidence suggesting that the fine-textured and coarse-textured morphotypes may differ in their salt tolerance. Furthermore, the study identified genotypic differences among the fine-textured accessions, although it did not explicitly examine their population structure. Because fine-textured accessions provide the primary germplasm for turf breeding, further investigation of them is warranted.

In this study we have characterized intraspecific variation in salt tolerance in fine-textured *P. vaginatum* as related to genotypic variation in this economically important morphotype. We assayed the shoot biomass and ionome profiles of 44 accessions representing 17 unique genotypes at multiple salinity treatments and performed population genetics analysis on a dataset that included an additional 7 genotypes to ask the following questions: 1) Does ionomic variation among genotypes correspond to genome-wide genetic differentiation (population structure)? 2) Do independently collected and maintained accessions of identical genotypes exhibit different ionomic profiles, indicating phenotypic plasticity for this trait? 3) Do tissue ion concentrations correlate with salt tolerance (measured as change in biomass between low and high salt treatments) for individual genotypes? 4) If so, to what extent are these correlations attributable to population level differences?

## METHODS

### Plant Material

Plants used in salinity tolerance phenotyping experiments were a subset of the fine-textured *P. vaginatum* accessions examined by Goad et al. (2021), including wild collections from throughout the southeastern US and USDA GRIN lines. A total of 44 accessions represented 17 unique genotypes (Table S1). An additional 7 genotypically unique accessions from Goad et al. (2021) were included in the population genetics analysis but were not phenotyped.

### Population genetic analyses

Raw genotyping-by-sequencing (GBS) genotype calls for the 24 unique fine-textured genotypes from Goad et al. (2021) were filtered by removing sites with MAF < 0.05 and missing data > 0.1. Population structure was then assessed by principal component analysis (PCA), performed in PLINK (Purcell et al. 2007), and assignment to genetic populations with ADMIXTURE (Alexander et al. 2009). For comparisons between populations, genotypes that were not unambiguously assigned to a single population in ADMIXTURE were placed into populations based on having ≥50% assignment to that population.

### Salt tolerance assays

Three stolon cuttings of each accession, consisting of a single node and connected shoot tissue, were transplanted to a 6 × 5 cm pot containing a clay growth medium (Turface Field and Fairway, Profile Products LLC, Buffalo, IL, USA) in the Donald Danforth Plant Science Center Plant Growth Facility. Plants were allowed to establish for 6 weeks and were watered twice daily to run-through with tap water. For the first week plants were covered with domes to prevent dehydration. For the next three weeks they were allowed to grow uncovered. They were then trimmed to 2 cm above the soil line and allowed to continue growing for the remaining two weeks.

Plants were then moved to one of three experimental flood trays where each tray contained one replicate of each accession with its position randomly assigned. A Raspberry Pi computer was programmed to flood all three trays simultaneously with nutrient solution from a shared reservoir twice daily to represent tidal inflows (Fig. S1). The pump flooding a given tray was automatically shut off once the solution reached a sensor placed 1 cm above the soil line; the solution then passively drained back into the reservoir over a 30 min time period. The nutrient solution consisted of 230 L of a 50/50 mix of tap water and a 2x concentration of Jack’s CA-MG 15-5-15 (JR Peters Inc., Allentown, PA, USA) with NaCl added according to treatment. Plants were allowed to acclimate to the flood tray for two days with no added NaCl before experimental treatments began, at which point they were trimmed to 2 cm above the soil line.

Plants in flood trays were exposed to increasing levels of salt over a period of 12 weeks with the salt level increased every two weeks after the initial acclimation period, for a total of six experimental treatments (salt levels) for each plant. Increase in the salinity of the nutrient solution was measured by the electrical conductivity of the solution (ECw, units dS/m) at two-week intervals. The ECw of each treatment and the amount of added NaCl were as follows: 2.5 dS/m = 0 g L^−1^, 10 dS/m = 6.9 g L^−1^, 20 dS/m = 13.8g L^−1^, 30 dS/m = 20.7 g L^−1^, 40 dS/m = 27.6 g L^−1^, 50 dS/m = 34.5 g L^−1^. The first week at each salinity level served as an acclimation period, after which the aboveground tissue of each plant was trimmed to 2 cm above the soil line and discarded. After the second week at the same salinity level plants were again trimmed to 2 cm; however, this time tissue was saved and dried for at least one week before being used in subsequent analyses.

To prevent increases in salinity concentration due to evaporation, the nutrient solution was changed weekly, and electrical conductivity was tested daily with a PINPOINT Salinity Monitor (American Marine, Ridgefield, CT, USA). The conductivity of the solution never exceeded the target value by more than 2.5 dS/m, so no adjustment was necessary over the course of a week. A sensor failure during the acclimation period of the 20 dS/m treatment resulted in an overflow of one flood tray. The remaining solution was no longer sufficient to fill all three trays so the nutrient solution was replaced with a fresh batch before the next watering event.

### Tissue weight measurements and ionomics sample preparation

Two weights were measured for each dried leaf tissue sample: the total mass of the tissue sample harvested from the plant (collected biomass) and the weight of the subset of that tissue used for ionomics analysis (sample weight). These two values were measured concurrently during ionomics sample preparation. The entirety of each dried sample was transferred to a glass tube and the weight was recorded as the collected biomass. The target weight for the ionomics pipeline was 60-125 mg; therefore, if the collected biomass for a sample was greater than 125 mg, tissue was removed from the tube until the sample weight was within the target range. For plants with less than 125 mg collected biomass, the measures for sample weight and collected biomass were identical. Some samples, particularly those in higher salt treatments, had collected biomass weights below the target sample range of 60 mg. Ionomics analysis was still performed on these samples, and the effect of low sample weight was controlled statistically as described below.

### Ionome measurements

Concentrations of B, Na, Mg, P, S, K, Ca, Mn, Fe, Co, Ni, Cu, Zn, As, Se, Rb, Mo, and Cd were measured with inductively coupled plasma mass spectrometry (ICP-MS) following the protocol of Ziegler et al. (2013). Briefly, each sample was digested overnight in 2.5 mL HNO_3_ containing 20 parts per billion (ppb) indium as an internal standard. Samples were then heated to 100° C and diluted to 10 mL with ultra-pure water containing yttrium as an internal standard. Concentrations for each ion were then measured on a Perkin Elmer Elan 6000 DRC-e mass spectrometer. Reported concentrations were corrected based on the indium and yttrium controls as well as a matrix matched control containing pooled samples from that run. This matrix match control was repeated after every 10^th^ sample to control for intra-run variation. Because the sample pool differed for each run, we statistically controlled for variation between runs by including the ICP run as an effect in our models as described below.

### Data filtering

Ionomics measurements were performed on samples from all salinity treatments; however, samples from the three highest salt treatments (30 dS/m, 40 dS/m and 50 dS/m) had excessive missing data due to dead plants and low sample weight (7 of 17 genotypes had 1 or fewer samples for the 30 dS/m treatment) (Table S2). This meant that sample sizes were inadequate to analyze ion concentration data for the three highest salinity treatments, and they were therefore excluded from subsequent analyses.

For the remaining 346 samples from the 2.5, 10, and 20 dS/m treatments, we applied a series of data filters. To reduce potential technical variation caused by samples with extremely low weights, we first removed ionomics samples that weighed less than 15 mg; this excluded 48 samples. Next, we identified samples with poor quality data across multiple ion concentrations. To do this we performed a PCA including the concentration of all ions for every sample. Five outliers were visually identified after plotting PC 1 and PC 2 and subsequently removed. For the remaining samples, we then identified outliers for each ion concentration within each salinity treatment following the algorithm from (Davies and Gather 1993) as implemented in the *outlierRemoveDataset* function in the *IonomicsUtils* package (https://github.com/gziegler/ionomicsUtils/). We used the default parameters which removes outliers surpassing a conservative 6.2 median absolute deviation (MAD). This final filtered dataset contained 293 samples (Table S3).

### Testing for correlations between weight and ion concentration

To test whether technical variation in elemental concentrations was due to sample weight variation, we performed another PCA on the cleaned dataset and used a linear regression to test for correlations between sample weight and the first two PCs. We also assessed whether variance in PCs could be explained by salinity concentration treatment or flood tray.

Because sample weight and collected biomass were not independent, we then tested whether sample weight correlations could instead be due to actual changes in tissue ion concentration rather than technical variation due to sample weight. We took the 144 samples that had greater than 60 mg collected biomass and ran a mixed-effects model for each ion concentration using the following formula:

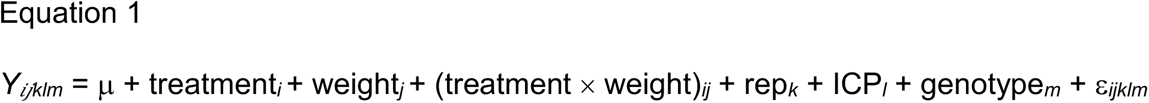

where *Y_ijklm_* is the tissue ion concentration; μ is the overall mean; treatment*_i_* is the *i*th salinity level; weight*_j_* is the collected biomass of the sample; (treatment × weight)*_ij_* is the effect of the interaction between sample weight and treatment; rep*_k_* is the flood tray as a random effect; ICP*_l_* is the random effect of the *l*th ICP-MS run; genotype*_m_* is the random effect of the *m*th genotype of the sample; and ε*_ijklm_* is the random error term.

We then ran the same model except with weight*_i_* as the effect of sample weight rather than collected biomass. If collected biomass was significant at a Bonferroni corrected threshold of *p* = 2.7 x 10^-3^ and sample weight was not, then the ion concentration was considered to be attributable to actual differences in ion accumulation between plants with different collected biomass rather than an artifact of sample weight.

### Genotype by treatment interactions

In order to test for significant interactions between genotype and treatment for each ion, we tested a model with the following equation for each ion concentration:

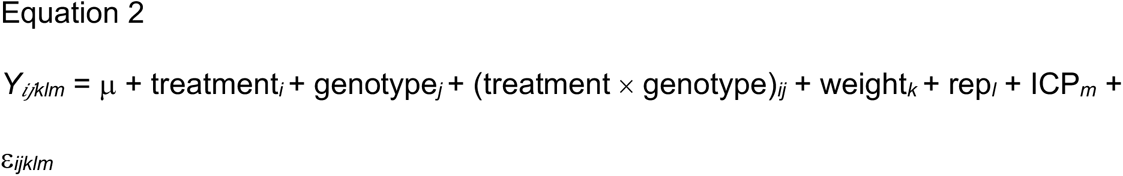

where *Y_ijklmn_* is the tissue ion concentration; μ is the overall mean; treatment*_i_* is the *i*th salinity level; genotype*_j_* is the random effect of the *j*th genotype; (treatment × genotype)*_ij_* is the effect of the interaction between genotype and treatment; weight*_k_* is the collected biomass of the sample; rep*_l_* is the random effect of the *l*th flood tray; ICP*_m_* is the random effect of the *m*th ICP-MS run; and ε*_ijklm_* is the random error term. We then checked for significance of the (treatment × genotype)*_ij_* term using a Bonferroni corrected threshold of *p* = 2.7 x 10^-3^.

### Within-genotype variation

To compare the level of variation between both genotypes and accessions within a genotype for each ion concentration, we tested the following nested mixed model:

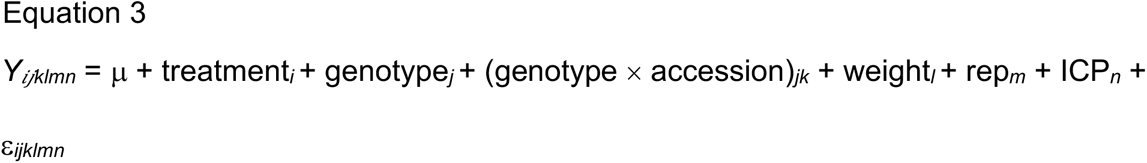

where *Y_ijklmn_* is the ion concentration; μ is the overall mean; treatment*_i_* is the *i*th salinity level; genotype*_j_* is the fixed effect of the *j*th genotype; (genotype × accession)*_jk_* is the nested effect of the *k*th accession within the *jth* genotype; weight*_l_* is the weight of the ionomics sample; rep*_m_* is the random effect of the *m*th flood tray; ICP*_n_* is the random effect of the *n*th ICP-MS run; and ε*_ijklmn_* is the random error term. We then checked for significance of the genotype*_j_* and (genotype × accession)*_jk_* terms using a Bonferroni corrected threshold of *p* = 2.7 x 10^-3^.

### Genotypic correlations between ion concentration and salt tolerance

To assay the level of salt tolerance of genotypes, we calculated the percent change in biomass due to increased salinity as the percent change in sampled biomass from the 2.5 dS/m treatment to the 30 dS/m treatment using the following equation: (biomass at 30 dS/m – biomass at 2.5 dS/m) / biomass at 2.5 dS/m. This value was calculated individually for each replicate of a genotype and then averaged. Biomass of samples that had been filtered out of the ionomics analyses due to low sample weight were included in this measure. If a plant had died by the 30 dS/m treatment, no percent change in biomass was calculated for it and that replicate was not included in the genotypic average. We tested for differences between the three genetic populations identified in our ADMIXTURE analysis for both collected biomass at 2.5 dS/m and the percent change in biomass using one-way ANOVAs.

For each ion we then calculated a best linear unbiased prediction (BLUP) for every genotype by extracting the genotype effect for each genotype from the following mixed model:

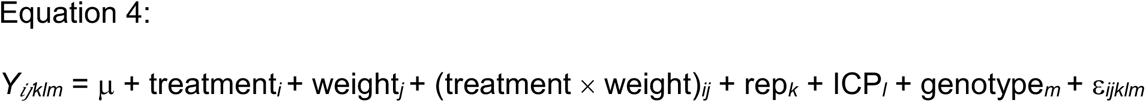

where *Y_ijklm_* is the adjusted ion concentration; μ is the overall mean; treatment*_i_* is the *i*th salinity level; weight*_j_* is the collected biomass of the sample; (treatment × weight)*_ij_* is the effect of the interaction between sample weight and treatment; rep*_k_* is the flood tray as a random effect; ICP*_l_* is the random effect of the *l*th ICP-MS run; genotype*_m_* is the random effect of the *m*th genotype of the sample; and ε*_ijklm_* is the random error term.

Since no genotype-by-treatment effects were significant with Equation 2, the interaction term was not included in the model. We then performed a PCA using the genotypic BLUPs as input. To test for genotypic correlations between the percent change in biomass and tissue ion concentration we performed a Spearman correlation between each BLUP or PC and the average change in biomass as calculated above. Correlations were considered significant with Bonferroni corrected threshold *of p* < 2.7 x 10^-3^.

## RESULTS

### Population structure

Population structure analysis of genotyping-by-sequencing (GBS) genotypes revealed three genetic populations in both the principal component analysis (PCA) and ADMIXTURE analysis (Fig. 1). The optimal K for the ADMIXTURE analysis as determined by the cross-validation (CV) error was 2; however, the CV error for K=3 was only slightly higher than K=2 and much lower than for all of the other potential K values (Fig. S2). We chose to use K=3 despite this slightly higher CV error because it better reflected groupings in both our genetic and phenotypic PCAs. One genetic population (labeled Population 1) was composed entirely of wild collections from the southeastern US Atlantic Coast and is the most genetically distant from the other two populations, particularly along PC 1 which explained 15.4% of the variance. PC 2 largely separated population 3 from the other two populations and explained 11.3% of the variance.

**Figure 1.**
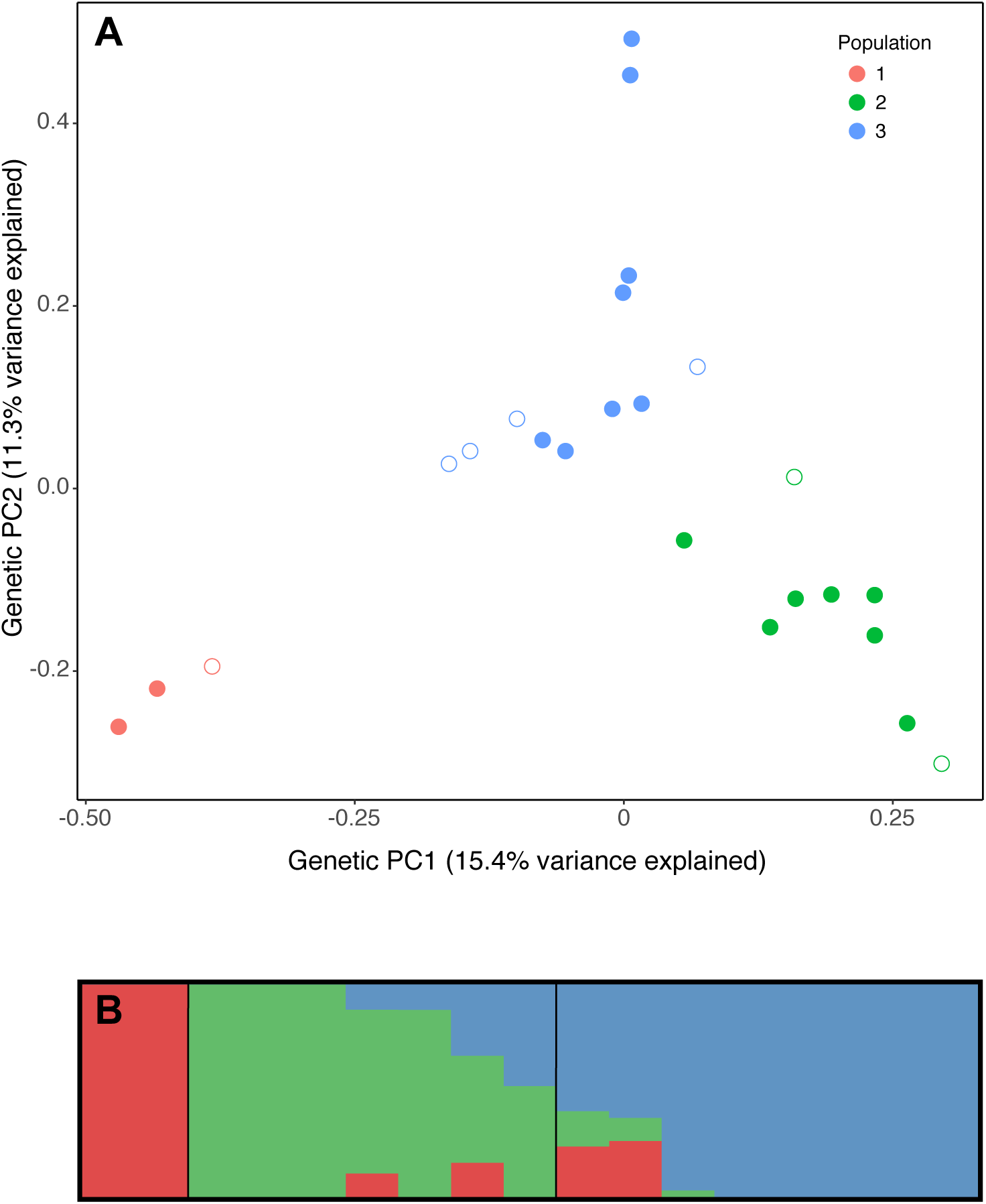
(**A**) A genetic PCA of all 24 unique fine-textured genotypes, based on data from Goad et al. (2020). The first and second principal components explain 15.4% and 11.3% of the variance respectively. Genotypes are colored based on assignment to genetic population by ADMIXTURE. Open circles indicate genotypes that were not phenotyped in this study. (**B)** A visualization of population assignment from ADMIXTURE for the subset of 17 genotypes for which we analyzed phenotype data. Vertical black lines divide genotypes into three populations (1, 2 and 3 from left to right). Admixed individuals were assigned to the population for which the membership coefficient was greater than 50%.

### Correlations between sample weight and ion concentration

To be sure that differences in ion concentration between samples were due to physiological differences and not technical error due to variation in sample weight, we performed a PCA on the samples from the 2.5, 10 and 20 dS/m treatments. The first two PCs (which explained 27.0% and 21.8% of the variance respectively) both correlated significantly with sample weight (PC 1, *r*^2^ = 0.14*, p* < 10^-11^; PC 2, *r*^2^ = 0.26*, p* < 10^-16^) (Fig. S3A). However, PC 2 was largely driven by salinity treatment (*r*^2^ = 0.62*, p* < 10^-16^) (Fig. S3B). These correlations indicate that sample weight may have a small effect on ionome quantification, but that it explains a much smaller proportion of the variance than treatment.

Since the weight of the ionomics sample and the total shoot biomass were sometimes the same (for samples up to 125 mg), we wanted to determine whether the correlations between sample weight and ion concentration may be caused by actual differences in accumulation between plants of different sizes rather than technical variation in sample weight. We examined the 144 samples within the target sample weight range (60-125 mg) because it included samples where collected biomass equaled sample weight. Within this range, no ions showed significant positive correlations between sample weight and ion concentration; however, concentrations of P, Rb and several heavy metals (Co, Ni, Cu, Zn and Cd) showed significant negative correlations between collected biomass and ion concentration after controlling for multiple hypothesis testing (*p* < 2.7 x 10^-3^) (Table S4). Based on this pattern, larger plants may have lower ion concentrations due to uptake being largely constant, resulting in the same quantity of ions being spread across more biomass and thus less concentrated. These results also suggest that the sample weight effect described above is largely driven by smaller samples (25-60 mg), as the sample weight effect is not significant when they are removed.

### Elemental differences within and between genotypes

Many genotypes were represented by accessions that were collected as cuttings directly from wild plants, so might have retained some residual within-genotype phenotypic variation reflecting environmental heterogeneity between collection locations. We found that only four elements (Co, Cu, Ni and Rb) varied significantly between accessions within a genotype and that all of the elements except S and Se had significant between-genotype differences (Table S4). Together these findings suggest that ionomic variation is largely stable but that plastic responses to environmental conditions may not entirely disappear when a plant is moved to a new environment.

### Elemental differences between populations

We quantified the overall elemental profiles of our genotypes by performing a PCA on the BLUPs generated for each of the 19 elements (Fig. 2A). The elemental profiles of genotypes clustered based on their population assignment. Furthermore, genetic PC 1 and elemental PC 1 were significantly correlated (Fig. 2B), as well as genetic PC 2 and elemental PC 3 (Fig. 2C), indicating that more distantly related genotypes have more dissimilar ionomic profiles. Population 1 (comprising two wild accessions from the southeastern US Atlantic Coast) appears to be particularly distinct from the other two populations both genetically and elementally.

**Figure 2.**
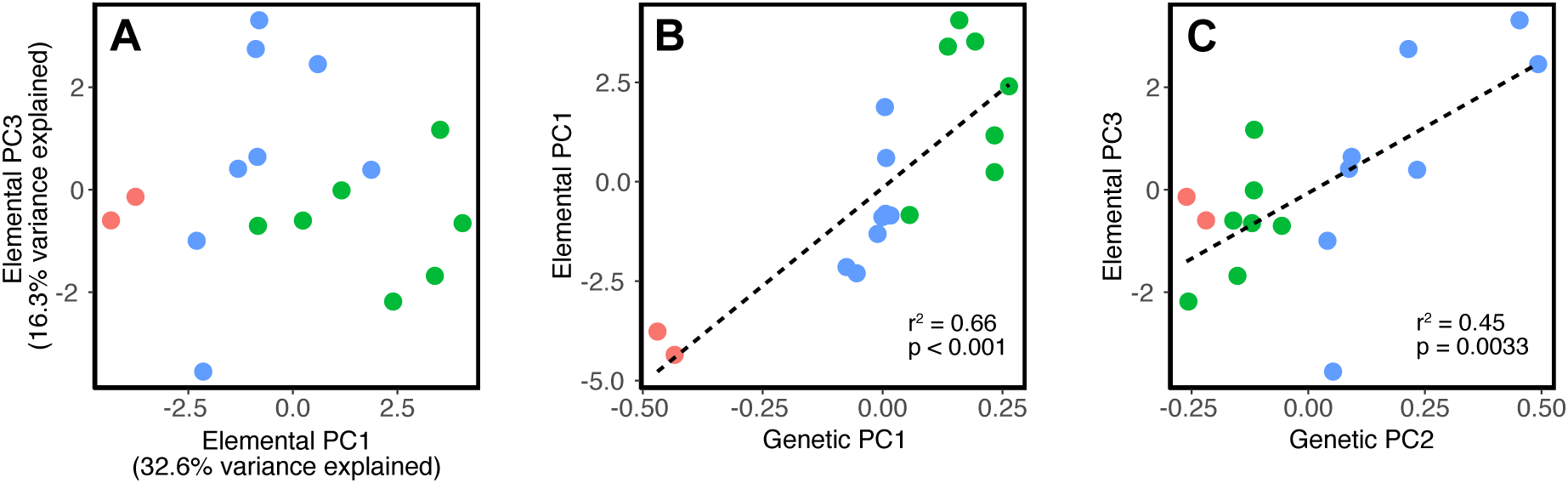
(**A)** PCA based on the BLUPs calculated for each ion concentration for each genotype. (**B)** The Spearman correlation between genetic PC1 and elemental PC1. (**C)** The Spearman correlation between genetic PC2 and elemental PC3. Genotypes are colored based on assignment to genetic population by ADMIXTURE.

### Biomass differences between populations

The three populations also significantly differed in both their overall biomass and the percent change in biomass at increased salt concentrations. During the initial no salt treatment (2.5 dS/m), the mean biomass for populations 1, 2, and 3 was 40.8, 120.1 and 94.9 mg, respectively (*F*(2,14) = 4.94, *p* < 0.05). All three populations showed reduced growth as the salt concentration increased and therefore had a negative percent change in biomass between the no salt and high salt treatments (2.5 dS/m and 30 dS/m). The magnitude of this change can be used as a measure of salt tolerance; notably, it differed significantly between populations (*F*(2,14) = 16.8, *p* < 0.001). Population 2 was least affected by the increased salinity with a -50.7% change in biomass. A larger reduction was observed in populations 1 and 3 which had similar percent changes in biomass of -78.1% and -80.8% respectively. Therefore, population 2 appears to be more salt tolerant than the other two populations.

### Correlations between ion concentration and percent change in biomass

After correction for multiple hypothesis testing, three separate elements showed significant correlations between the ion concentration BLUPs and percent change in biomass of individual genotypes. K and Fe showed significant positive associations (*r*^2^ = 0.55, p < 2.7 x 10^-3^ and *r*^2^ = 0.54, *p* < 2.7 x 10^-3^ respectively) while Ca showed a significant negative association (*r*^2^ = 0.50, *p* < 2.7 x 10^-3^) (Fig. 3, Table S5). This result suggests that shoot concentrations of these three elements may play some role in salt tolerance. No genotype-by-treatment interactions were significant, indicating that genotypes all adjust ion accumulation in the same way with increased salinity (Table S4). Consistent with our findings that different genetic populations also differ in their elemental profiles, these trends follow the expected patterns based on population structure. Genotypes from Population 2 maintain a higher relative biomass in increased salinity and have the highest K and Fe concentrations as well as the lowest Ca concentrations. Population 1 was on the opposite end of the spectrum for these measurements and Population 3 was intermediate. This pattern indicates that differential accumulation of these three ions between populations may reflect local adaptation for varying salinity levels in the wild.

**Figure 3.**
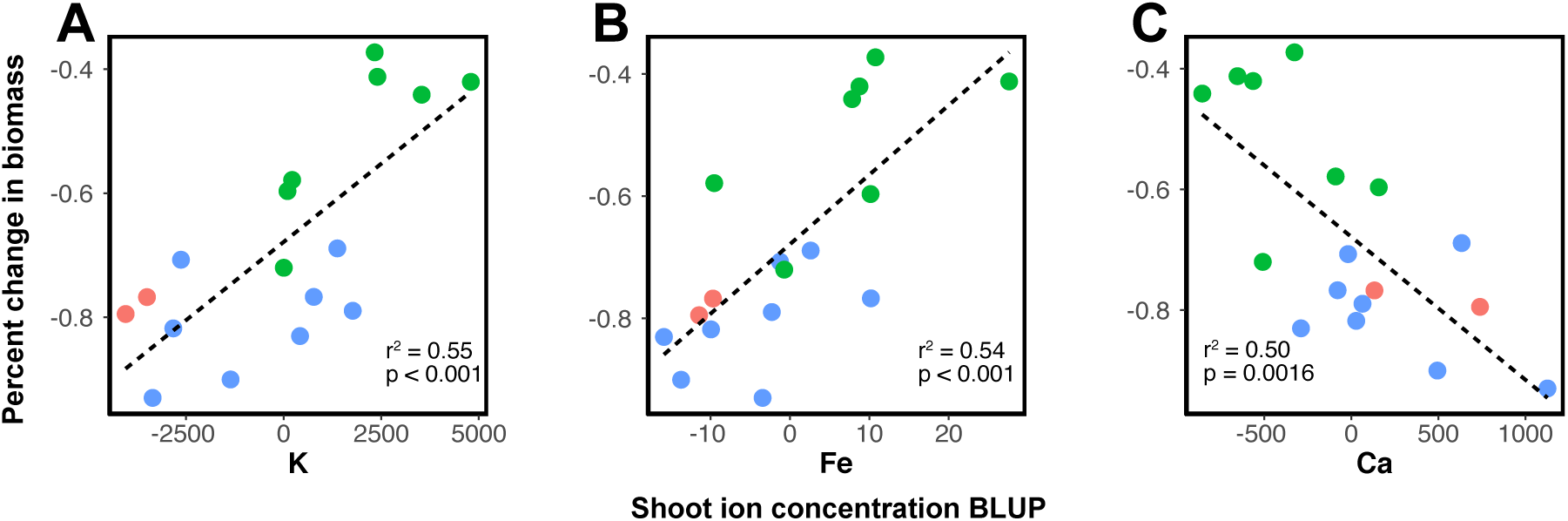
Correlations between the percent change in biomass at increased salinity and shoot concentration of K (**A**); Fe (**B**) and Ca (**C**). Genotypes are colored based on assignment to genetic population by ADMIXTURE.

## DISCUSSION

### Genetic and elemental profile variation within and between genotypes

Understanding salt tolerance variation in wild species requires consideration of population structure as well as variation in ionomic profiles among genotypes. Our population genetics analyses are the first to explore population structure within fine-textured *P. vaginatum* using genome wide SNP markers. We recovered both of the genetic populations previously identified in this species (Eudy et al. 2017) (corresponding to our Populations 2 and 3), in addition to a new third population which was unique to our collections from the US Atlantic Coast (Population 1) (Fig. 1). Our analysis of the elemental profiles of each genotype largely recapitulated the pattern seen in the population structure analysis, with strong correlations between the first PC of both the genetic and elemental datasets as well as between genetic PC 2 and elemental PC 3 (Fig. 2). This finding suggests that genotypic differences in elemental accumulation traits have diverged between populations and may be heritable. It is not clear whether this divergence is adaptive since none of the genetic or elemental PCs were significantly associated with variation in the percent change in biomass at increased salinity. It is possible that these population level differences could reflect adaptations for other stressful conditions that we did not test for, such as drought or heavy metal contamination.

We also found variation within genotypes for shoot accumulation of Co, Cu, Ni and Rb (Table S4). All of the accessions used in this experiment were taken as cuttings from wild plants or USDA germplasm rather than grown from seed. Therefore, it is possible that phenotypic plasticity or epigenetic differences between otherwise genetically identical plants from different locations could still be affecting phenotypes in our common garden experiment (Bossdorf et al. 2008). *Paspalum vaginatum* often propagates asexually in the wild (Goad et al. 2021) and plasticity in elemental accumulation could allow clones of a single genotype to have increased fitness across a wider variety of environments (Vrijenhoek and Parker 2009). It is also possible that within genotype variation could be due to rare polymorphisms that arose from somatic mutation within clonal lineages that were not detected in our GBS dataset. Whole genome sequencing including a focus on epigenetic modification (e.g. bisulfite sequencing) will be required to understand the relative role of somatic mutations and epigenetics in explaining the variation seen within genotypes.

### Correlations between elemental accumulation and salt tolerance

Maintaining tissue ion homeostasis is one of the major challenges faced by plants in saline environments. The hypertonic environment can cause dehydration unless the plant can regulate osmotic pressure by accumulating ions or organic osmolytes. In some cases Na is accumulated at a higher rate to maintain this balance (Baxter et al. 2010). This is not an ideal solution, however, as Na is cytotoxic at high concentration and can compete with the accumulation of other important ions. Therefore, tightly regulating the uptake of Na is vital for halophyte survival, and variation between genotypes in these mechanisms could be involved in differential salt tolerance. Like earlier studies (Lee et. al 2007; Guo et. al 2016) we did not find a significant correlation between salt tolerance, as measured by percent change in biomass, and Na accumulation within *P. vaginatum* (Table S5). We did, however, detect significant correlations for K, Fe and Ca. It is thus possible that differential accumulation of these three elements in the shoots could play a role in the variation in tolerance seen within fine-textured *P. vaginatum.* Below we discuss the potential biological roles that these three elements could be playing in salt tolerance variation.

### Potassium

K accumulation plays a key role in salt tolerance in both halophytes and glycophytes (Shabala and Cuin 2008). Due to its similarity to Na in both size and charge, K accumulation faces direct competition with Na. Compared to Na, however, K is more vital for plant function and less toxic (Shabala and Cuin 2008). Halophytic species tend to accumulate more K under saline conditions than glycophytes (Ghars et al. 2008; Edelist et al. 2009), and intraspecific variation in K accumulation is associated with increased salt tolerance in many glycophytes (Chen et al. 2005; Chakraborty et al. 2016). The role of K accumulation in salt tolerance variation within halophytic species is less well understood. Fine-textured *P. vaginatum* has been shown to have increased K accumulation compared to the less tolerant *P. distichum* and coarse-textured *P. vaginatum* accessions (Spiekerman and Devos 2020). This difference may be due to the increased size of adaxial leaf papillae in fine-textured plants; these structures sequester Na and K, which could allow for greater ion accumulation (Spiekerman and Devos 2020). The role that variation in K accumulation plays within fine-textured accessions is less well characterized. Previous studies have identified a difference in shoot K accumulation between the most and least tolerant accessions; however, there was no correlation between K and salt tolerance when comparing all of the sampled accessions (Lee et al. 2007; Guo et al. 2016; Wu et al. 2020). Our results are thus the first to show that K accumulation is associated with increased tolerance across multiple fine-textured varieties. It is possible that, as with leaf morphology differences between coarse- and fine-textured morphotypes, variation among fine-textured plants in the size and function of the adaxial leaf papillae could explain this trend. Since we measured accumulation in the entire shoot, we could not differentiate between the K content of the adaxial leaf papilla and other structures. A more thorough survey of the adaxial papillae in a large sample of fine-textured genotypes would be useful to test this hypothesis on their role in K and Na accumulation.

### Iron

The accumulation of Fe was also associated with increased tolerance in our experiment. This finding is consistent with observations in glycophytes that increases in environmental Fe concentration, and thus accumulation, can alleviate the negative impacts of high salinity (Ghasemi et al. 2014). Fe plays an important role in photosynthesis and homeostasis of reactive oxygen species. In plant tissues, free Fe is sequestered in a non-toxic form by storage proteins called ferritins. Halophytes exposed to salinity have been shown to have increased expression of ferritin genes and to exhibit higher Fe concentrations, particularly in the chloroplast where ferritin may improve the efficiency of photosynthesis during salt stress (Paramonova et al. 2004; Jithesh et al. 2006). Within *P. vaginatum*, the Fe transporter protein IRT1 has been shown to be differentially expressed in high salinity environments, and transgenic yeast containing a copy of the *IRT1* gene from *P. vaginatum* showed increased salt tolerance (Chen et al. 2016). Together these findings suggest that Fe accumulation, ferritin production and IRT1 may have some influence in determining the level of salt tolerance of a *P. vaginatum* genotype.

### Calcium

Calcium plays an important role in plant physiology particularly as a structural component of the cell wall and membranes and as a secondary messenger in a wide range of signaling networks (Thor 2019). It is also involved in plant salt tolerance as part of the SOS pathway which regulates K and Na homeostasis (Mahajan et al. 2008). In multiple species, supplemental Ca has been shown to ameliorate the negative effects of salinity stress (Carvajal et al. 2000; Sun et al. 2009; Yousuf et al. 2015; Larbi et al. 2020). Therefore, our finding that increased shoot Ca concentrations are associated with reduced salt tolerance in *P. vaginatum* was surprising. It is possible that the shoot Ca concentration may not directly affect salt tolerance, but instead reflects differences in transpiration and water conservation between genotypes.

Unlike many solutes, long range transport of Ca to the shoots primarily follows an apoplastic pathway and is therefore strongly tied to the rate of transpiration rather than membrane bound calcium transporters which are primarily involved in signaling on a more local scale (White 2001). Within *P. vaginatum* some genotypes have been shown to more quickly reduce transpiration in response to drying soil as a water conservation strategy (Johnson et al. 2009). It is thus possible that in our experiment the drying of the growth medium between flooding events could have triggered these responses. If this is the case, the plants that maintained normal transpiration rates for a longer period of time could have accumulated more shoot Ca, but been more susceptible to osmotic stress at higher salinity concentration due to increased water loss. A more focused study involving explicit measures of transpiration rate will be required to better understand the interactions between salt tolerance, transpiration and Ca accumulation in *P. vaginatum*.

## Conclusions

Intraspecific variation in salt tolerance within halophytes remains poorly understood. Here we showed that genetic populations of *P. vaginatum* differ in their elemental accumulation traits and that salt tolerance is associated with the accumulation of specific ions. This finding suggests that a population level approach to studying halophytes may be a valuable, and so far, untapped, source of knowledge regarding salt tolerance genes and mechanisms for breeding more tolerant crop varieties. Breeders interested in improving seashore paspalum for use as a turf grass may be able to select for optimal K, Fe and Ca accumulation traits to improve salt tolerance. Other elements which showed differential accumulation between genotypes and populations may also play a role in adaptation to other stressors (e.g. drought, heavy metal contamination, poor soil quality) and could also be of interest for turf breeding.

## ACKNOWLEDGMENTS

We thank Kevin Riley, Sally Fabbri and the rest of the Danforth Center greenhouse staff for assistance with plant care and irrigation system development; Marshall Wedger and Taylor Aubuchon-Elder for assisting with tissue collection; Melissa Jurkowski for running the ionomics pipeline; and Greg Ziegler for providing barcoded sample labels and data analysis advice. Funding for this project was provided by the US Golf Association (project 2016-35-605). DMG was supported by the William H. Danforth Plant Science Fellowship and the Donald Danforth Plant Science Center.

## DATA AVAILABILITY

Demultiplex and trimmed reads from Goad et al. (2021) are available at the NCBI SRA under project ID PRJNA669382. The SNP dataset from Goad et al. (2021) are available on Dryad at https://doi.org/10.5061/dryad.vt4b8gtqh. Ionomics and biomass data are included in Table S3. All scripts used for data analysis and for controlling the irrigation system are available at https://github.com/david-goad/paspalum_ionomics.

## SUPPLEMENTAL MATERIAL

Supplemental tables submitted as separate excel file.

Table S1. Sample information including unique genotype ID, population assignment, and collection location for samples from Goad et al. (2021).

Table S2. Number of replicates of each genotype at each sat treatment before and after data quality filters were applied.

Table S3. Sample information including IDs, run information, inclusion in final analysis, biomass measurements, sample weight, and raw ionomics data

Table S4. Associations between tissue ion concentration and terms from Equation 1 and 2. Bold text indicates significance at the Bonferroni corrected threshold of *p* < 0.0027.

Table S5. P values and correlation coefficients for the association between change in biomass at increased salinity and shoot ion concentrations or principle components. Bold text indicates significance at a Bonferroni corrected threshold of *p* < 0.0027.

**Figure S1.**
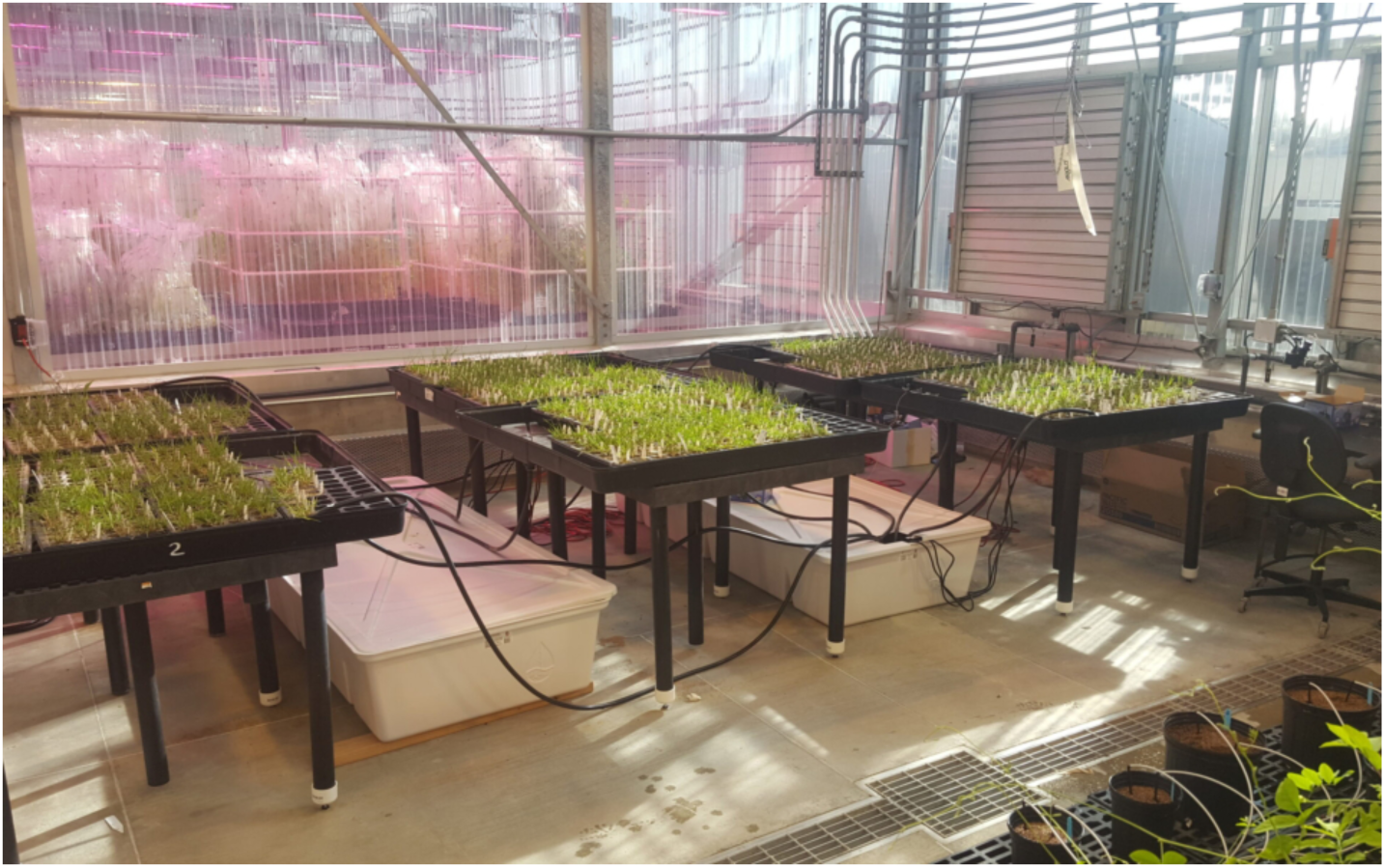
Flood tray design. Three of the six flood trays pictured (one from each table) were used for this experiment. Each tray contained 1 cloned replicate of each named accession. All three trays were flooded from and drained into the same reservoir for the duration of the experiment.

**Figure S2.**
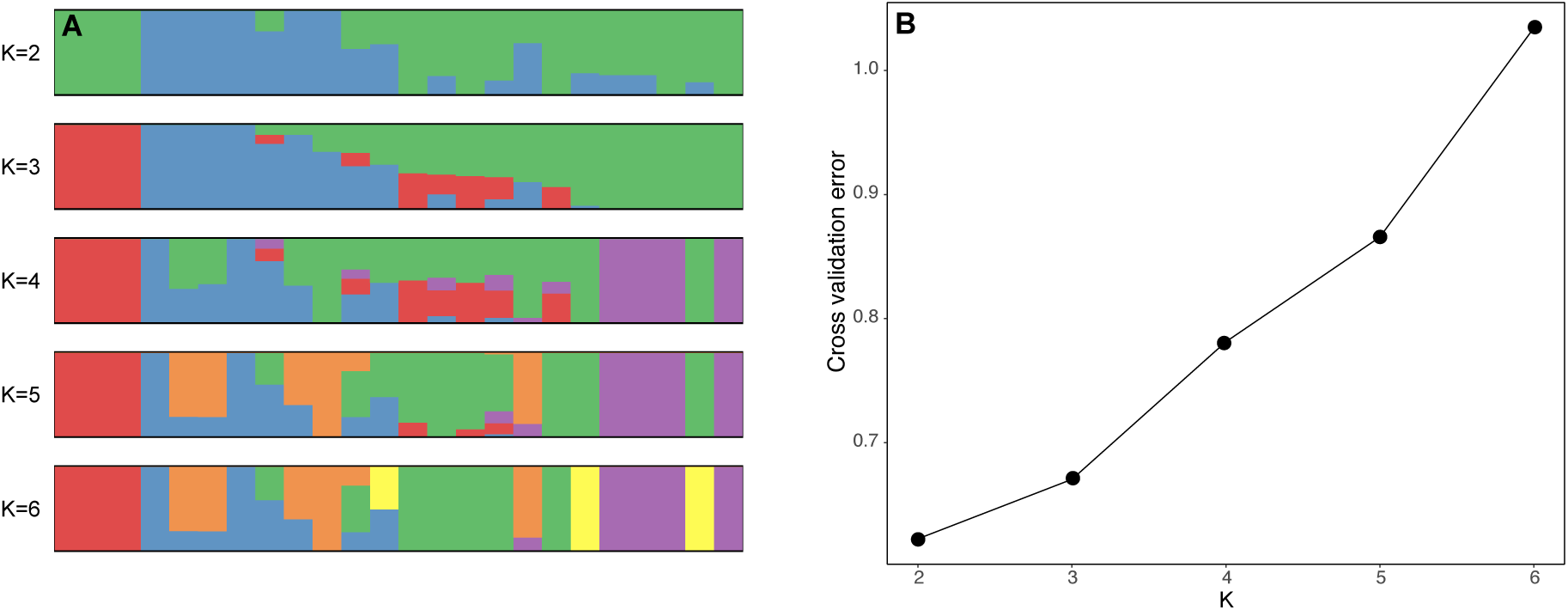
(**A)** ADMIXTURE Plots for K values 2 to 6. (**B**) Cross validation error for ADMIXTURE runs with K values from 2 to 6.

**Figure S3.**
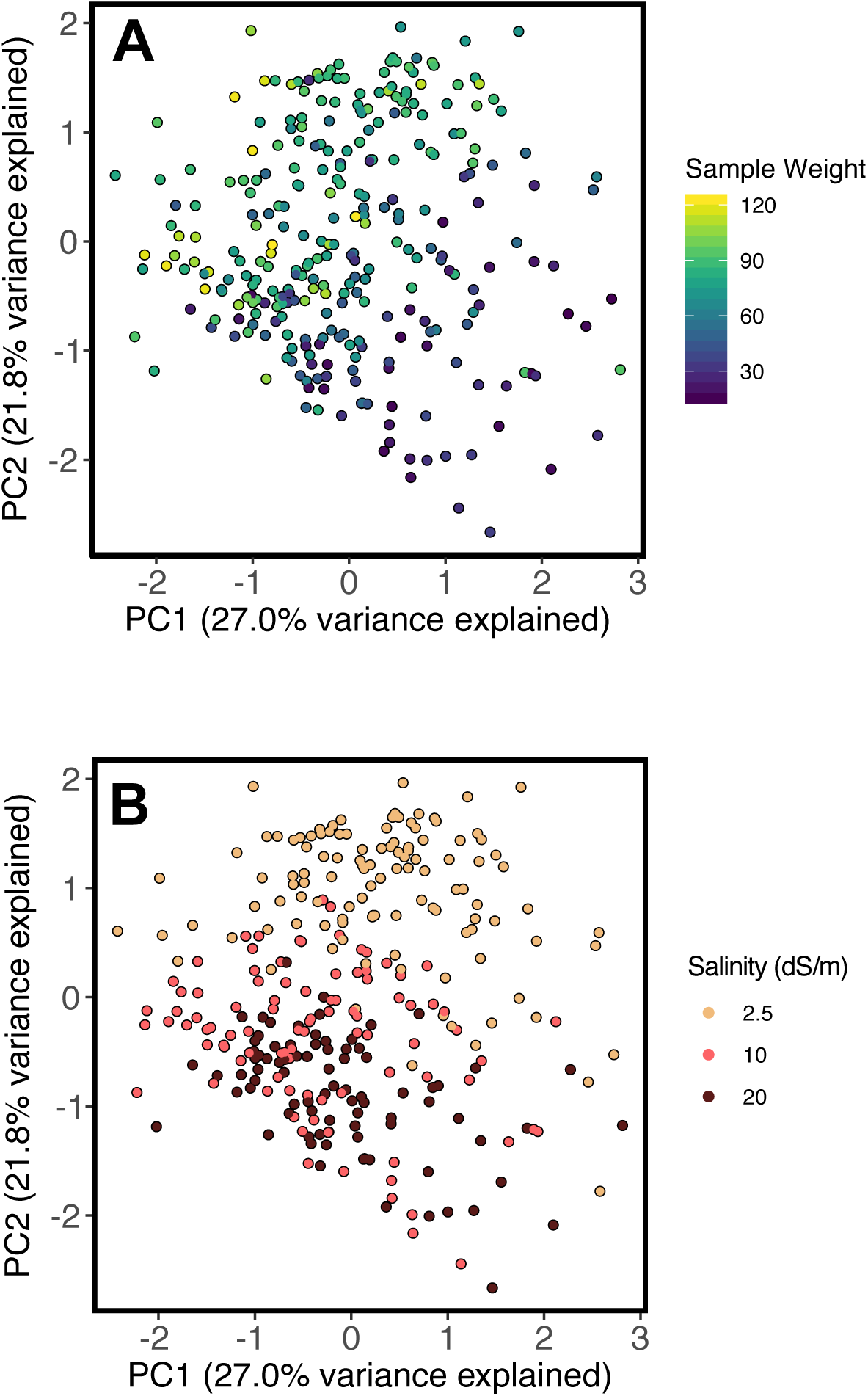
PCA of ionome profiles for all post-filtering ionomics samples in the 2.5, 10 and 20 dS/M treatments. Samples are colored by sample weight (**A**) and treatment (**B**).

## Notes

### Competing Interest Statement

The authors have declared no competing interest.

